# Tumor-Antagonizing Fibroblasts Secrete Prolargin as Tumor Suppressor in Hepatocellular Carcinoma

**DOI:** 10.1101/2021.01.04.425182

**Authors:** Barbara Chiavarina, Roberto Ronca, Yukihiro Otaka, Roger Bryan Sutton, Sara Rezzola, Takehiko Yokobori, Paola Chiodelli, Regis Souche, Antonio Maraver, Gavino Faa, Tetsunari Oyama, Stephanie Gofflot, Akeila Bellahcène, Olivier Detry, Philippe Delvenne, Vincent Castronovo, Masahiko Nishiyama, Andrei Turtoi

## Abstract

Systemic treatment of hepatocellular carcinoma (HCC) targets the tumor microenvironment (TME) by combining immunotherapy with angiogenesis inhibitors. Cancer-associated fibroblasts (CAF) are important components of the TME, however their targeting in clinics remains challenging. To this end, the existence of tumor supressing CAF and their poor characterisation is a major conundrum. Starting from proteomics and single cell analysis, we outline CAF heterogeneity in HCC and describe a subtype of CAF that express a novel tumor-related protein prolargin. Upon secretion prolargin is deposited in the TME where its levels positively correlate with patient outcome (HR=0.37; p=0.01). *In vivo*, tumors with lower prolargin expression display faster progression (5-fold; p=0.01) and stronger angiogenesis. Mechanistically, aggressive HCC cells degrade prolargin using matrix metalloprotease 3 (MMP3). We show that prolargin binds and inhibits several growth factors, key to tumor progression. Inhibiting prolargin degradation combined with sorafenib, demonstrated superior tumor control compared to sorafenib treatment alone. In conclusion, prolargin-expressing CAF have tumor-antagonizing properties. Stabilizing prolargin tumoral levels should be considered for systemic therapy of HCC, involving CAF in existing TME targeting.

## Introduction

Hepatocellular carcinoma (HCC) is the third leading cause of mortality from cancer worldwide.^1, 2^ By 2035, HCC is expected to become the second cause of cancer-related deaths (1.3 million people per year; GLOBOCAN, WHO). Despite the rising number of patients, HCC treatment remains limited, with patients facing poor survival outcomes (5-year, 18%).^3^ Surgery and liver transplantation give the best curative chances (5-year survival rates of up to 70%), while molecular therapies were until now only palliative in nature. Indeed sorafenib, an RTK inhibitor and standard of care for HCC for the past decade, prolongs survival of non-operable HCC patients for 3 months when compared to placebo.^4^ Only until recently, no treatment managed to outperform sorafenib. Indeed, newest clinical trials showed that combination of atezolizumab (anti-PD-L1) and bevacizumab (anti-VEGF) lead to a significant improvement of patient outcome compared to sorafenib. ^5^ The rationale of combining the VEGF blockade with immune checkpoint inhibitor stems from VEGF pro-angiogenic as well as immunosuppressive activity in promoting tumor progression.^6^ Independently, angiogenesis per se has been well identified as a major driving force in HCC ^7^ and the efficacy of the previous first line treatment (sorafenib) rested mainly with its ability to impair this process.^4^ HCC represents therefore a unique example where harnessing multiple components of the tumor microenvironment proved more beneficial than targeting cancer cells themselves.

Tumor microenvironment is a complex entity that harbours many cellular subtypes, of which immune cells and CAF are in majority.^8^ While tumor-infiltrating immune cells are well established as a heterogeneous population with both cancer suppressing and promoting functions, CAF have been, and still largely are, regarded as genuine cancer cell collaborators.^9^ In a static picture of a tumor, CAF engage in a multitude of tumor supporting processes.^10^ This accounts for numerous attempts to target and deplete CAF in tumors in the hope of achieving a therapeutic benefit. However, beyond animal studies, where these strategies have indeed shown to be successful,^11, 12^ we have yet to clinically prove the expected benefit in humans.^13^ Current data clearly show that indiscriminate CAF targeting accelerates tumor progression.^14, 15^ Similar to the concept of immunity, fibroblasts are naturally programmed to suppress rather than to support cancer growth.^16-18^ Carcinogenesis is a dynamic process, thus fibroblasts, along with other stromal cells, have to undergo programming steps in order to shift the balance from tumor protection to promotion.^19, 20^

In line with the thought that not all CAF are the same, recent studies have demonstrated the co-existence of several CAF subtypes within the same tumor. Some of the subtypes are characterized with immunosuppressive and chemoresistant phenotypes.^21-23^ However, we have had very little insight into the population of CAF that have tumor antagonizing properties. This is probably due to the fact that this cell population is in minority in advanced tumors. Only a handful of studies have characterized molecular markers of tumor-antagonizing CAF.^24-26^ Recently, we uncovered asporin as a CAF-derived tumor suppressor protein in triple negative breast cancer that inhibits TGF-β1 signaling.^27^

In this study, we use for the first time a combination of innovative proteomics approach with single cell sequencing data analysis to better characterize the CAF in human HCC. Among numerous stromal proteins already mainly characterized as tumor promoters, a newly identified cancer-related protein, prolargin (PRELP), was found to be specifically expressed by portal-derived CAF. Here, we examine the molecular function of PRELP by highlighting its novel ability to antagonize tumor progression (*in vitro* and *in vivo*) and propose its significance for conceiving future HCC treatments.

## Materials and Methods

Full description of Experimental Procedures is found in the Supplemental Material section.

### Patient information and clinical samples

The study was conducted in compliance with the Declaration of Helsinki. The use of human material was approved by the institutional ethical committees of the University Hospital of Liège and Gunma University Graduate School of Medicine. Clinical information concerning the patients used in the present study are outlined in the Supplemental Data, **Table S1**.

### Single Cell RNA-seq Analysis

We reanalysed previously published single cell RNA-seq analysis involving 9 HCC patients.^28^ Processed 10X Genomics (Pleasanton, CA, USA) data were downloaded from GEO repository (GSE125449). They were imported into R computational environment (4.0) and then processed using *Seurat 3*.*1* package and default parameters.^29^ CAF identification/origin (stellate or portal fibroblast) was inferred by comparing the present data set to the single cell study of Dobie et al.^30^ Ligand-receptor analysis was conducted using *SingleCellSignalR* package.^31^

### Immunohistochemistry (IHC) & Immunofluorescence (IF)

Formalin-fixed paraffin-embedded (FFPE) tissue sections were prepared from primary hepatocellular carcinoma lesions and from xenografted tumors. Tissue samples were sliced from paraffin blocks (5-μm sections), deparaffinated three times in xylene for 5 min and hydrated in a methanol gradient (100%, 95%, 70%, and 50%). Blocking of unspecific peroxidase activity was performed for 30 min with 3% H2O2 and 90% methanol. Diluted ImmunoSaver solution (1:200 in DDW, Nisshin EM, Tokyo, Japan) was used for antigen retrieval. Tissues were treated with Protein Block Serum-Free solution (Protein Block Serum-Free Ready-to-Use, catalog no. X0909, Dako, Glostrup, Denmark) for 30 min at room temperature, preceding the primary antibody incubation. Primary antibodies used for IHC and IF are described in the Supplemental Materials section. They were detected with corresponding secondary antibody: i) donkey anti-sheep IgG H&L (HRP polymer) (catalog no. ab214884, Abcam) or ii) M-Histofine Simple Stain MAX PO (Multi) (catalog no. 414152F, Nichirei Biosciences Inc., Tokyo, Japan). For IHC signals were detected using 3,3’-diaminobenzidine tetrachlorhydrate dihydrate (DAB) in 5% H2O2. The slides were counterstained with hematoxylin and dehydrated with ethanol and xylene for mounting. For multiplexed immunohistochemistry signals were detected using Opal™ 4-Plex Kit according to the manufacturer’s protocol (catalog no. NEL810001KT; Perkin Elmer, Waltham, MA, USA). For more details see Supplemental Materials section.

### Murine in vivo models

All experimental procedures used in the current work were performed in accordance with the ARRIVE ethical guidelines.^32^ They were reviewed and approved by the Ethical Committees of the Bioresource Center in Gunma University (Japan) and the French National Committee of animal experimentation. To evaluate the impact of fibroblast-secreted PRELP on tumor development, HUH7-RFP-Luc cells (0.5×106 cells) and NLF-shNT or -shPRELP (0.5×106 cells) were suspended in 20 µl of cell culture medium and growth-factor-reduced Matrigel (catalog no. 356230; BD Biosciences) 1:1 and co-injected into the liver of six-week old athymic nude mice (n=5 per condition).

## Results

### Prolargin is a novel CAF-derived protein of human HCC

We have previously developed a proteomic method to characterize extracellular and membrane-bound proteins in fresh human tumors.^33, 34^ Here we employed this methodology for the first time on fresh human HCC and non-tumoral liver specimens from 6 individual patients (**Figure S1**). We identified 144 significantly modulated proteins in at least 3 out of 6 patients (**Figure S1a**). The proteins in question mainly participate in extracellular matrix organization (**Figure S1b**). Pathway analysis (KEGG) revealed alterations in ECM-receptor interaction, focal adhesion and notably the PI3K-Akt signalling pathway (**Figure S1b**). These intriguing findings led us to perform further network analysis using STRING software. The most prominent network was observed around a strong collagen cluster, complemented by known collagen-interactors such as fibronectin (FN1), periostin (POSTN) and tenascin (TNC) (**Figure S1c**). We also observed asporin (ASPN), a small leucine-rich proteoglycan (SLRP) that was previously described by our group as a fibroblast-derived tumor suppressor in triple negative breast cancer. The analysis highlighted another SLRP protein; prolargin (PRELP), which to the best of our knowledge has not yet been functionally described in cancer. We thus sought to better understand the significance of prolargin modulation in the context of human HCC. To this end, we first aimed to delineate the cell population responsible for prolargin expression. We have re-analysed recently published single cell RNAseq data in human HCC.^28^ As outlined in **Figure 1a**, 17 cell populations were identified using tSNE analysis, including 3 distinct CAF cell populations. Comparative analysis with recently published data on liver fibrosis,^30^ clearly identified these clusters as hepatic stellate cell-derived CAF (CAF_HSC), vascular smooth muscle cell-derived CAF (CAF_VSMC) and portal fibroblasts-derived CAF (CAF_Port). Based on relevant markers permitting to distinguish these three CAF subtypes (**Figure S2a)**, prolargin mRNA was exclusively found in portal fibroblasts-derived CAF (**Figure 1b**). Examination of PRELP protein expression and localisation in normal liver confirmed its localisation in portal areas and not in the liver sinusoids (**Figure 1c**). PRELP was found expressed in the tumor (α-SMA^+^/PRELP^+^ cells), however not all CAF were positive for PRELP (α-SMA^+^/PRELP^-^), emphasizing the expected heterogeneity from single cell analysis (**Figure 1d**).

**Figure 1:**
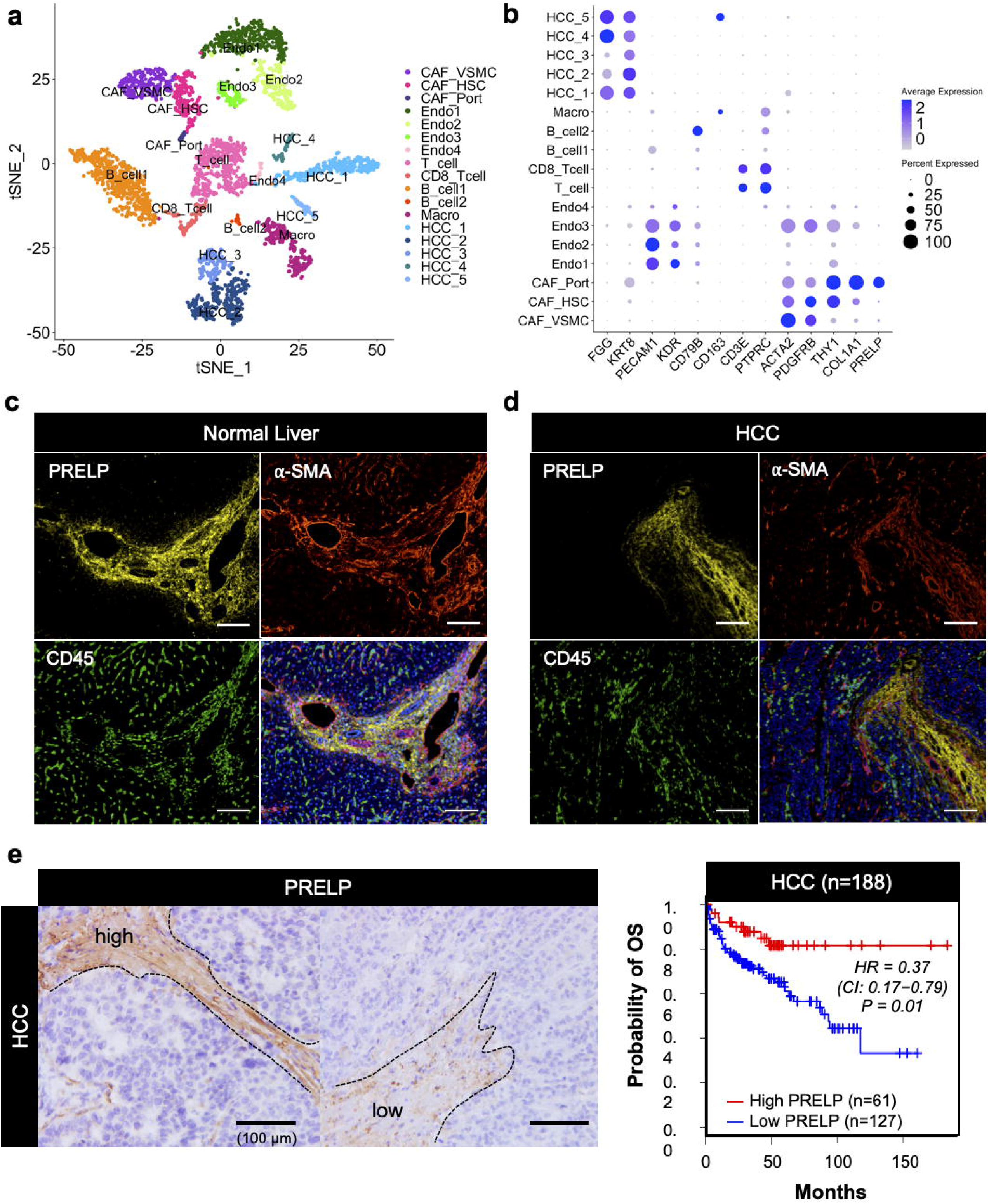
Prolargin is expressed by CAF deriving from portal fibroblasts. **a**. t-SNE plot of different cell populations found in the HCC from 9 patients. **b**. PRELP expression in individual cell populations. Panels **a** and **b**: single cell RNAseq analysis of previously published dataset GSE125449. **c**.**/d**. IF analysis of PRELP expression in human normal liver (**c**) and liver cancer (HCC) PRELP is co-stained with αSMA (stellate/fibroblast cells) and CD45 (immune cells). Shown are representative images of 20 individual cases. **e**. IHC analysis of PRELP expression in human HCC (N=188 cases). Shown are representative cases with high/low PRELP expression in the HCC lesion. High PRELP levels correlate with good clinical outcome in human HCC. Kaplan-Meier survival curve indicates the probability of overall survival (OS).

Based on single cell data, CAF_Port represented only a minority population of all CAF in HCC, which were mainly derived from stellate and smooth muscle vascular cells (**Figure 1a**). Thus, we next sought to estimate the importance of the individual CAF populations with respect to their communication with other stromal and cancer cells in HCC. For this purpose we examined ligand-receptor expression on individual populations and used inference of intercellular networks to quantify and score these interactions.^31^ Surprisingly, and as shown in the **Figure S2b**, CAF_Port although in minority had 3-times more interactions with other HCC cell populations compared to CAF_HSC or CAF_VSMC.

Having identified the cells that express prolargin, we next sought to better understand its function in HCC. To this end, we performed IHC analysis on HCC patients with the objective to examine its relationship with patient survival (clinical details are outlined in **Table S1**). Our findings showed that prolargin levels positively correlated with good overall survival (HR = 0.37, p = 0.01, **Figure 1e**).

### Extracellular prolargin levels are controlled by MMP-mediated degradation

In order to functionally study prolargin we established a cell line model. This model was based on a set of five HCC cell lines that vary in their aggressiveness, as well as one primary cell line of normal hepatic fibroblasts (denoted NLF). In clonogenicity assay, HLE, HLF and Alexander cells formed large and dense colonies, while HepG2 and HUH7 in this regard were less aggressive (**Figure 2a**). In accordance with their aggressiveness, on the molecular level HepG2 and HUH7 cells had a pronounced epithelial phenotype (CDH1^high^/VIM^low^), while Alexander, HLE and HLF were mesenchymal-like (CDH1^low^/VIM^high^) (**Figure S3a**). Proteomic analysis of conditioned media (CM) derived from all the five cancer cell lines suggested that HepG2 and HUH7 cells were particularly well differentiated, with certain secretory aspects similar to hepatocytes (e.g. secretion of albumin) (**Figure S3b**). In contrast to this, we found increasing levels of several proteases in CM from HLE, HLF and Alexander compared to HepG2 and HuH7 cells (**Figure 2b**).

**Figure 2:**
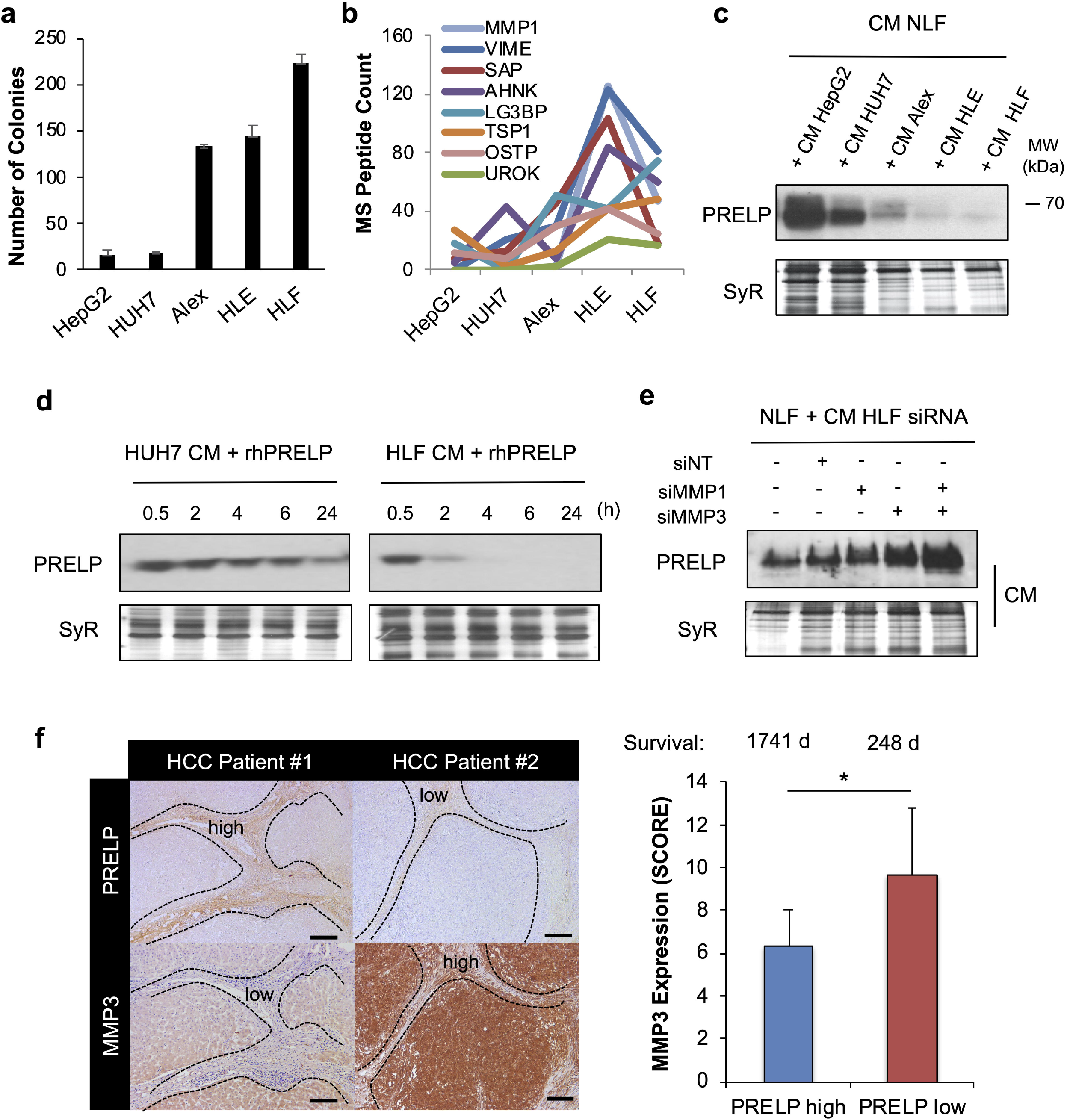
Extracellular prolargin levels depend on cancer cell MMP1 and MMP3 expression. **a**. Colony formation ability of 5-cell HCC panel. **b**. Proteomic analysis of the secretome obtained different HCC cells. Displayed are the proteins that increase in expression along with increasing aggressiveness of HCC cells lines. **c**. Levels of extracellular NLF-derived PRELP following the challenge of NLF with conditioned media (CM) from cancer cells. **d**. CM from HUH7 and HLF cells incubated with recombinant human PRELP at 37°C. **e**. Treatment of NLF with CM from HLF cells obtained following their silencing for MMP1 and MMP3. Panels **c, d** and **e**: Total protein stain with SYPRO Ruby (SyR) were used for normalization. **f**. Expression of MMP3 in a selection of PRELP high- and low-expressing HCC cases (20 patients each). Stroma is delineated with dashed lines; shown are serial sections. Average survivals are indicated in days (top of the graph). *denotes statistical significance with p<0.05, error bars are standard deviations of means.

In line with the evidence from single cell analysis, prolargin expression was not detectable in any of the cancer cells, while it was positive in NLF (**Figure S4a**). Next, we assessed the response of NLF following 48h treatment with CM from the individual cancer cell lines. As prolargin is a secreted protein, we sought to verify the modulation of prolargin levels in the medium of CM-treated fibroblasts (intracellular prolargin levels were unaffected, **Figure S4a**). Extracellular levels revealed that prolargin was gradually degraded upon NLF treatment with media of cancer cells with increasing aggressiveness potential HepG2 < HUH7 < Alexander < HLE < HLF (**Figure 2c**). To delineate the mechanism of prolargin degradation we incubated CM from HUH7 and HLF cells with recombinant prolargin and evaluated the respective protein levels at different time points. In remarkable contrast to HUH7 cells, CM obtained from HLF cells was able to rapidly degrade exogenously added prolargin (**Figure 2d**). To further narrow down which proteases are involved we tested two MMP inhibitors, batimastat and GM6001, as well as amiloride HCL (inhibitor of urokinase-type plasminogen activator (uPA)), and a broad protease inhibitor cocktail (Roche Complete). As demonstrated in **Figure S4b**, both batimastat and GM6001 were able to inhibit prolargin degradation in the CM from HLF cells, suggesting that MMP play a major role. We next sought to determine the identity of the MMP involved, and thus performed a pull-down of recombinant prolargin incubated with different cancer cell-CM. Mass spectrometry analysis revealed MMP1 and MMP3 proteins to be the potential degradation enzymes (**Figure S4c**). The selective silencing of MMP1 and MMP3 (as well as the combination of both, **Figure S4d**) confirmed MMP3 as the main protease involved in the cancer cell-mediated degradation of secreted prolargin (**Figure 2e** and **Figure S4e**). Based on these findings we next examined human HCC for MMP3 and prolargin expression on a limited cohort of patients (N=40). The analysis showed a significant inverse relationship between the levels of the two proteins, suggesting that the mechanism described *in vitro* might also apply *in vivo* (**Figure 2f**).

### Prolargin is a CAF-derived tumor suppressor with antiangiogenic features

Based on our observations made in both patients and *in vitr*o, we assumed that prolargin might function as a fibroblast-derived tumor suppressor. In order to verify this, we grafted human HCC cells with fibroblasts expressing different levels of prolargin in the liver of nude mice. Human NLF were silenced for prolargin expression using shRNA (**Figure S5a/b**). These cells, or their non-silenced counterparts, were then grafted with HUH7 cancer cells. A luciferase expression vector in HUH7 cells permitted a regular monitoring of tumor development *in vivo*. Our findings show that HUH7 cells, when co-injected with prolargin-depleted fibroblasts, developed faster growing HCC in comparison to the same cells co-injected with fibroblasts that were not silenced for prolargin (**Figure 3a**). To further evaluate the mechanism behind this tumor suppressive behaviour of prolargin, we analysed the tumors recovered from HUH7-NLF orthotopic co-injection. Initial histological inspection revealed clear signs of necrosis in the prolargin proficient tumors. Prolargin deficient tumors, were compact with no sign of necrosis. These observations suggested that prolargin proficient tumors were less oxygenated. We further explored this by evaluating the density of vasculature/endothelial cells (CD31) in both tumors (**Figure 3b**). In contrast to those tumors where prolargin was silenced, the vasculature in wild-type tumors presented higher number of smaller vessels typically found during neo-vascularization. These findings collectively prompted us to put forward the idea that CAF-derived prolargin could be interfering in the crosstalk between cancer and endothelial cells, leading to the suppression of tumor angiogenesis. To explore this, we used ligand-receptor analysis on the single cell data. We specifically charted the interaction of portal fibroblast-derived CAF which express prolargin with different sub-clusters of endothelial cells identified in the t-SNE analysis (see **Figure 1a**). Portal fibroblast-derived CAF engaged in numerous strong interactions with all endothelial cell clusters (**Figure 3c** and **Figure S5c**), notably via collagen binding to vascular endothelial growth factor receptors 2 and 3 (VEGFR2/3), decorin interaction with c-MET and angiopoietin-like 4 (ANGPTL4) interaction with tyrosine kinase with immunoglobulin-like and EGF-like domains 1 (TIE1). To further test the relationship between prolargin, tumor angiogenesis and endothelial cells we performed proliferation and spheroid sprouting assays using human umbilical vein endothelial cells (HUVEC). CM obtained from Alexander or HLE cells when mixed with recombinant prolargin were significantly less potent in inducing both HUVEC proliferation and sprouting (**Figure 3d**). Together these data suggested that prolargin exerts its activity through a possible interaction with pro-angiogenic growth factors in the extracellular space.

**Figure 3:**
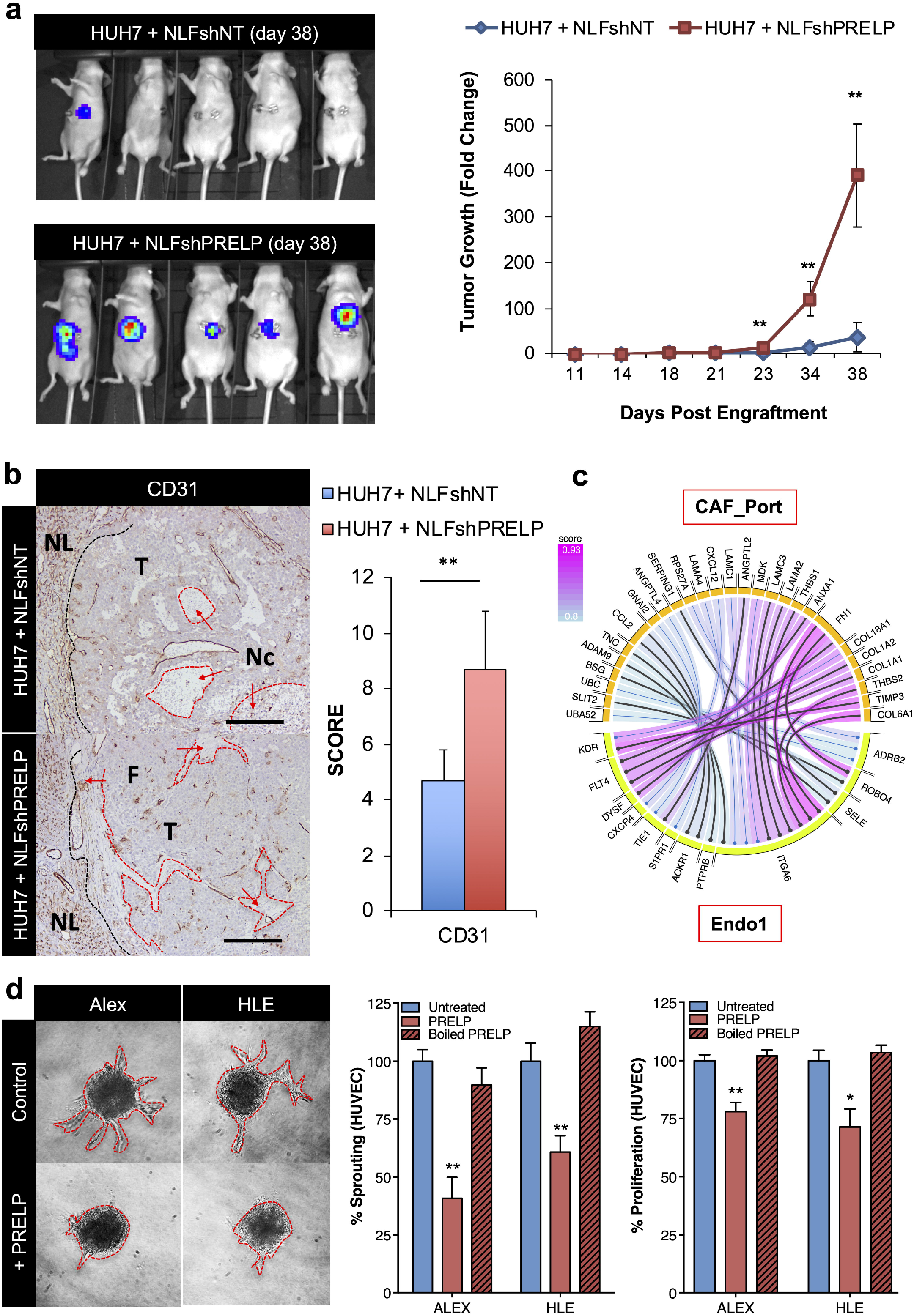
Prolargin inhibits tumor growth by suppressing angiogenesis. **a**. (*left panel*) Bioluminescence imaging of control and prolargin silenced orthotopic xenografts. Red color indicates the highest and blue the lowest fluorescent intensity. (*right panel*) Fold-changes of average radiance in liver tumors (compared to day 7 post engraftment). The data are presented as mean ± standard error of means (N=5 for each group). Statistical significance was calculated using Student’s *t*-test (*: 0.05<p **: p<0.01). **b**. Histological analysis of control and PRELP silenced tumors, showing the distribution/quantification of murine vasculature/endothelial cells (CD31). Labelled are: normal liver (NL), tumor region (T), fibrosis (F) and necrosis (Nc). **c**. Ligand-receptor analysis based on single cell RNAseq data. Shown is the interaction between portal-fibroblast derived CAF and endothelial cell cluster 1 (see **Figure 1** and **Figure S5**). **d**. Proliferation and sprouting assay using HUVEC treated with conditioned media of Alexander and HLE cells supplemented with recombinant PRELP. **b** and **d**: */**denote statistical significance with p<0.05 and p<0.01 respectively, error bars are standard deviations of means.

### Prolargin binds several key growth factors and inhibits their activity

In order to investigate the mechanism behind the prolargin anti-tumor function, we performed surface plasmon resonance. Prolargin was immobilized on the chip and 15 pro-angiogenic growth factors were screened for binding (**Figure 4a**). Of the 15 growth factors, prolargin demonstrated binding affinity towards 7 molecules, most notably FGF1, FGF2, HGF and TGF-β1. Interestingly, no binding was detected for VEGFA or VEGFC, a prominent angiogenic/lymphangiogenic factors. We next sought to understand how prolargin could interact with these growth factors on molecular level. To do so, we studied the ligand-receptor interactions using protein docking methods. In absence of crystal structure, we constructed a homology model of PRELP (**Figure S6a/b**) and used it to dock with known structures of several growth factors. As shown in the **Figure 6a/b**, PRELP model adopted a solenoid (horseshoe) folding. The most obvious structural feature of PRELP that could affect ligand binding is shape complementarity (Sc).^35^ The computed ‘Sc’ ratio was highest between PRELP and HGF (Sc∼0.7) and lowest for the FGF1 ligand (Sc∼0.6). Further, each of the high-affinity ligands (FGF1/2 and HGF) in this study docked within the curve of the solenoid, while TNFA and VEGFA did not (**Figure 4b**). In support of this, previous studies using other leucine-rich repeat (LRR) containing proteins have demonstrated similar docking behaviour when the interaction between LRR and growth factors takes place.^36^ We thus concluded that high-affinity binding in these growth factors likely favour productive binding that maximizes surface area exposure between receptor and ligand, as well as maximizing H-bond interactions between PRELP and the various growth factors. Lower affinity growth factor interactions, for example between PRELP and FGF1, are predicted to bind with a smaller footprint on the inner surface of PRELP.

**Figure 4:**
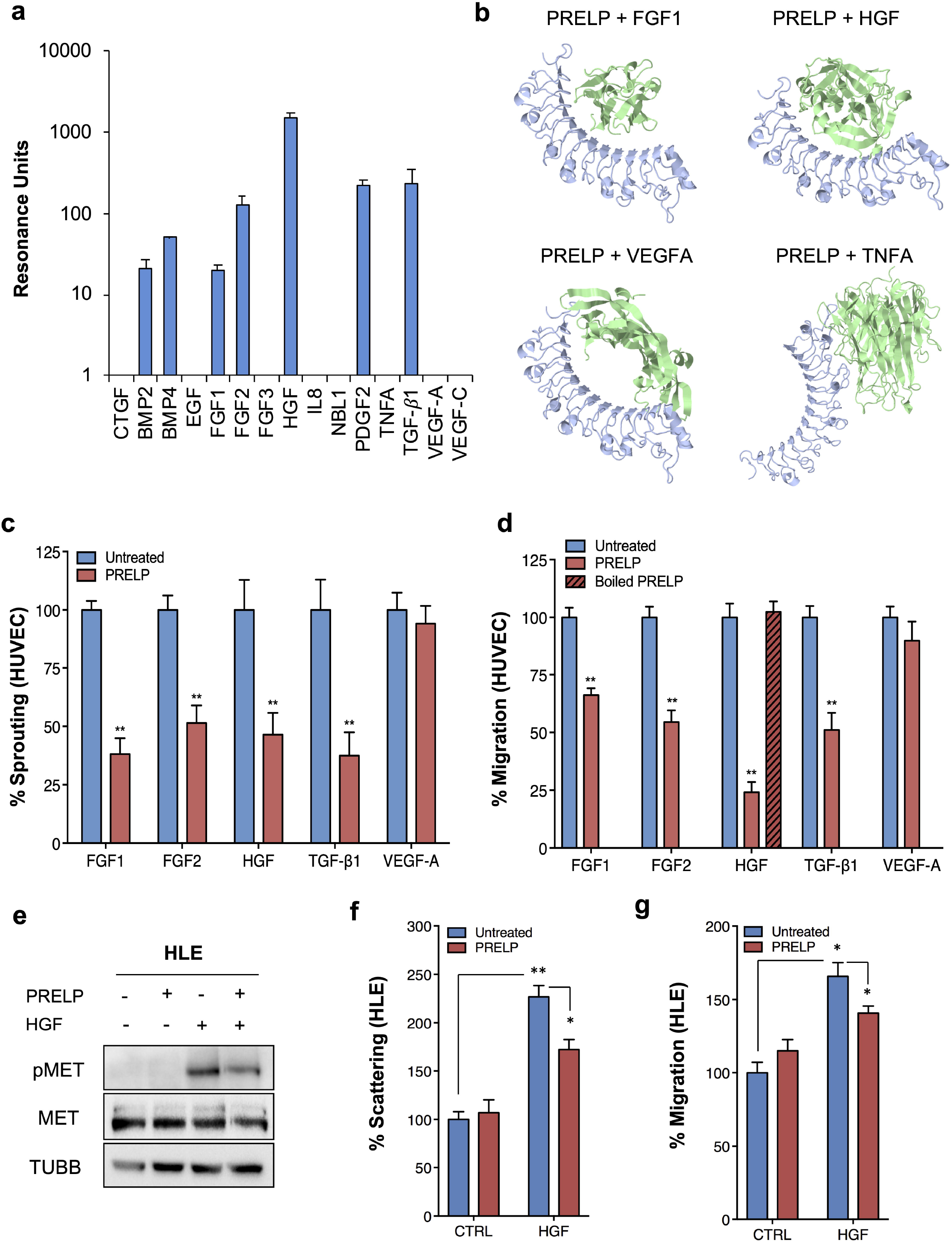
Prolargin binds pro-angiogenic growth factors and inhibits their activity. **a**. SPR profiling of growth factor binding to recombinant PRELP. **b**. Highest scoring docking solutions between PRELP and FGF1, HGF, VEGFA and TNFA. **c**. Sprouting assay using HUVEC treated with various growth factors or growth factors pre-incubated with recombinant prolargin. **d**. Migration assay with HUVEC; same conditions as panel (**c**). **e**. Western blot analysis of MET receptor activation following the treatment of HLE cells with PRELP (P), HFG (H) or a mix of PRELP and HGF (H+P). TUBB is used as loading control. **f**. Scattering assay with HLE cells treated with HGF or HGF pre-incubated with prolargin. **g**. Migration assay with HLE cells. Panels **a, c, d, f** and **g** */** denote statistical significance with p<0.05 and p<0.01 respectively, error bars are standard deviations of means.

Next, we sought to examine if prolargin binding neutralizes or enhances the activity of the respective growth factors. We thus incubated recombinant prolargin with FGF1, FGF2, HGF, TGF-β1 and VEGF respectively, and then tested growth factor activity on HUVEC as target cells (sprouting and migration assays) (**Figure 4c/d**). Prolargin inhibited the function of all the tested growth factors, with exception of VEGF, whereas prolargin denaturation by high temperature suppressed its functional activity (shown for HGF in **Figure 4d**). Among the different growth factors, HGF is particularly relevant in the context of HCC. The latter is known for its key ability to promote angiogenesis, fuel therapy resistance and enhance metastasis. Incubation of prolargin with HGF inhibited the activation of c-MET in both Alexander and HLE cells, as well as HUVEC (**Figure 4e** and **Figure S6c**). This inhibition functionally resulted in reduced scattering and migratory capacity of HLE cancer cells (**Figure 4f/g**). Together the data suggested that prolargin’s ability to bind and inhibit a broad range of growth factors has multiple paracrine effects, both on cancer and endothelial cells.

### Inhibition of prolargin degradation and VEGFR targeting is a meaningful therapeutic combination

Considering that prolargin binds to multiple growth factors and inhibits angiogenesis we aimed to test if stabilization of prolargin using MMP inhibitors (such as batimastat) could be therapeutically exploited. Knowing however that prolargin has no effect on VEGFA/VEGFC, and hence the activity of their receptors, we sought to combine prolargin stabilization with another treatment that should act mainly on VEGFR-1/-2/-3. To assess this we selected sorafenib as this inhibitor acts mainly on VEGFR-1, VEGFR-2, VEGFR-3 and PDGFR-β.^37^ Additionally, sorafenib is already used for systemic treatment for HCC.^38^ To reduce the number of animals used in subsequent experiments we examined the impact of individual drugs and combination on HLF tumors in chick chorioallantoic membrane model (CAM). We chose HLF cells because they were the most aggressive in our panel and we have added to them NLF cells as natural source of prolargin. CAM HCC tumors were treated with placebo, sorafenib, batimastat or a combination of sorafenib and batimastat. Both drugs were used in the nM range, known to be sufficient to inhibit only MMP and VEGFR activity, without causing toxicity to cancer cells. As shown in **Figure 5a**, within the 7 days of tumor development we could not observe a significant effect with either drug alone. However, the combination of sorafenib and batimastat significantly reduced the tumor volume (50% reduction) compared to placebo or either treatments alone. Western blot quantification of prolargin expression in CAM tumors confirmed higher levels following batimastat treatment (**Figure 5b**). Based on these data we proceeded to the *in vivo* study of mice bearing orthotopically transplanted HLF cells that were treated with two lines of therapy: i) sorafenib and ii) sorafenib/ batimastat combination treatment. As shown in **Figure 5c**, HLF tumors started to develop at day 38 despite the on-going treatment with sorafenib, suggesting that *in vivo* these cells can rapidly develop tolerance to the dose used. Remarkably, the combination of sorafenib and batimastat successfully controlled tumor growth, with no significant tumor growth until the end of the experiment (day 52).

**Figure 5:**
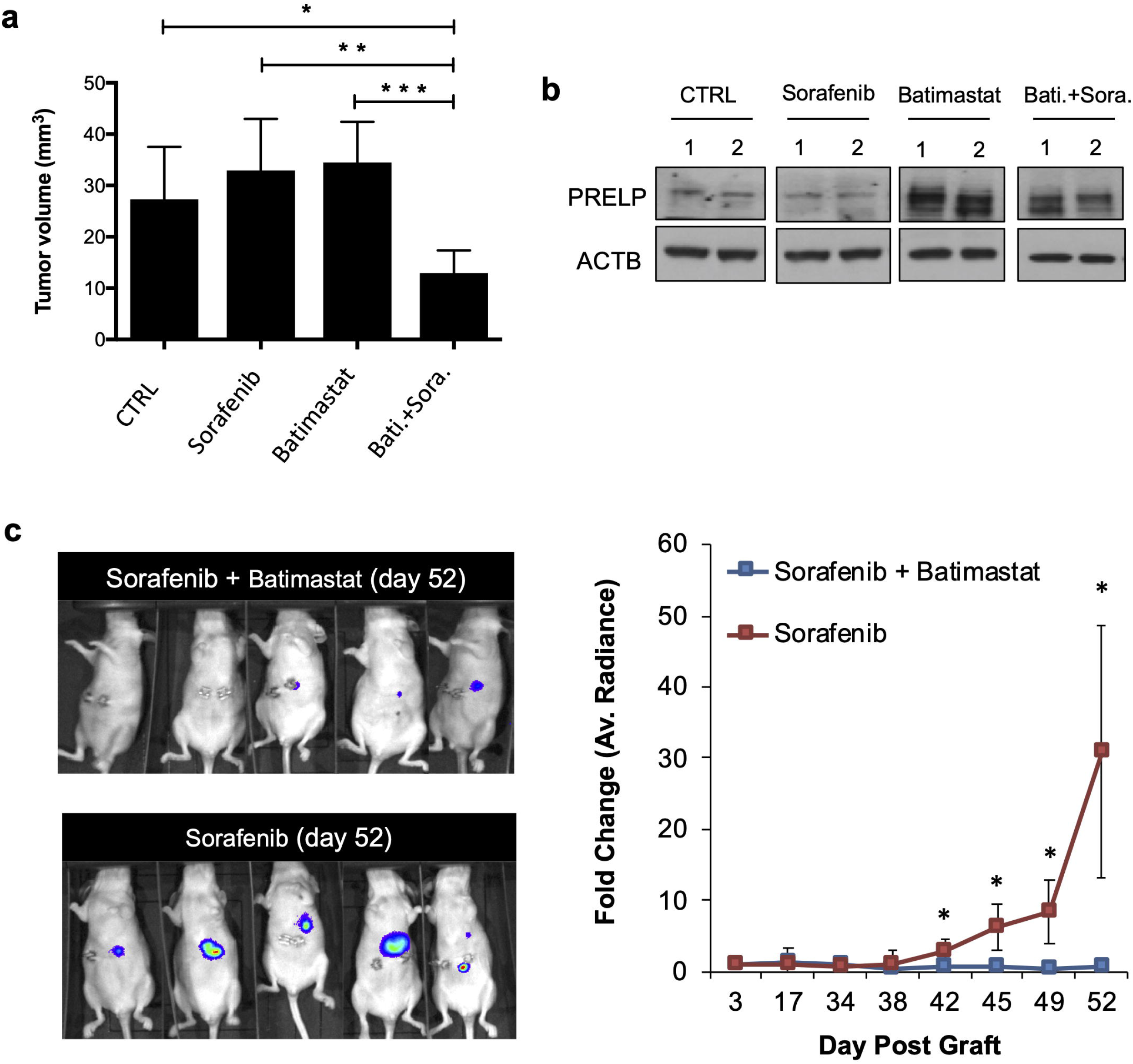
Combination of MMP inhibitor batimastat with sorafenib results in strong tumor control. **a**. Combined treatment with batimastat and sorafenib, reduces tumor growth on CAM. HLF cells were co-injected with NLF and then treated with placebo, sorafenib and batimastat alone or in combination. After 7 days, tumors were collected and tumor volume was measured. */**/ *** denote statistical significance with p<0.05, p<0.01 and p<0.001 respectively, error bars are standard error of means; N=5 for each group. **b**. WB analysis of PRELP levels in CAM-derived tumors following their treatment with batimastat/sorafenib. **c**. (*left panel*) Bioluminescence imaging post orthotopic engraftment of HLF tumor cells. Red color indicates the highest and blue the lowest fluorescent intensity. (*right panel*) Fold-change of average radiance in liver tumors (compared to day 3 post engraftment). The data are presented as mean ± standard error of means (N=5 for each group). Statistical significance was calculated using Student’s *t*-test (*: 0.05<p).

## Discussion

Cancer-associated fibroblasts have traditionally been regarded as a supportive part of a tumor microenvironment. Contrary to this view, here we describe prolargin, a novel protein characterizing a tumor-antagonizing subtype of CAF in human hepatocellular carcinoma. We identify a CAF-derived protein with a pleiotropic capacity to bind and inhibit the function of several crucial growth factors. This in turn leads to a significant decrease in HCC development *in vivo*, owing at least in part to the inhibition of tumor angiogenesis. Whether alterations of prolargin expression are also causative for HCC development still remains to be determined. Our data however suggest that prolargin probably belongs to a larger group of stromal proteins that need to be lost or their levels diminished for successful progression of HCC. Until recently, tumor-suppressive functions of CAF were not widely acknowledged, and thus the observations of both tumor promoting and suppressive roles were regarded as contrasting and controversial.^11, 14, 15, 39^ This apparent dichotomy could be reconciled by understanding that CAF are a heterogeneous population. One direct reason for CAF heterogeneity is their respective cells of origin. CAF can be derived from multiple cellular sources.^40^ In the HCC, there are at least three obvious source of CAF: stellate cells, smooth muscle vascular cells and portal fibroblasts. In the current study, we confirmed this and showed based on single cell RNAseq data those three populations of CAF in a collection HCC cases. Importantly here, prolargin was solely expressed by portal fibroblast-derived CAF, and it was this population (despite being in minority) that had the most numerous ligand-receptor interactions with other cell types in HCC. CAF heterogeneity has been recently shown in several types of tumors. Interestingly, in a recent study, Öhlund and collaborators21 identified two CAF populations in pancreatic ductal carcinoma (PDAC), one with high α-SMA and the other with low α-SMA expression. Costa et al.^22^ have shown the existence of four CAF subtypes in breast cancer and Li et al.^41^ identified two distinct CAF subtypes in colorectal cancer. While the functional significance of different CAF population is not always clear, a few studies succeeded in identifying certain CAF subpopulations that bear particular tumor promoting functions. For example, Su et al.^23^ identified CD10^+^GPR77^+^ CAF as a relevant population promoting chemoresistance and stemness in lung cancer. Costa et al. identified CD29^+^FAP^+^ CAF as key to recruiting CD4^+^CD25^+^ immunosuppressive T cells in breast cancer. Interestingly, Su et al. showed that the percentage of CD10^+^GPR77^+^ CAF in both breast and lung tumors is prognostic of disease-free survival. Conversely, findings from a study by Costa et al. lacked a similar correlation concerning CD29^+^FAP^+^ CAF. This raises a pertinent question if these newly identified CAF subpopulations do bear clinical significance. One evident explanation could be the discrepancy between gene and protein expression and the difficulty to predict functions from gene expression data (such as single cell RNAseq). To make things more complex, and as the present case of prolargin shows, intra- and extra-cellular levels of secreted proteins can additionally vary due to other regulation mechanisms such as degradation. Our present data based on protein expression clearly shows that the levels of prolargin secreting fibroblasts are positively correlated with good clinical outcome in HCC patients (overall survival). Admittedly, the patient cohort used in the present study is not representative of all HCC patients. This is because they include specimens exclusively from operated cases (30% were in addition also transplanted), and hence these patients already have better survival than non-operable patients. Nevertheless, the ability of prolargin to distinguish patients with better overall survival in operated HCC patients, highlights its strength as a prognostic marker.

Successful HCC treatment already involves targeting the microenvironment/stroma while in the future certainly also some aspects of personalized medicine. For every new treatment combination, clinicians will require markers that enable them to better guide the therapy. One of the most important findings of our study is the identification of prolargin as a prognostic and potential therapeutic biomarker. Indeed, we show that prolargin degradation directly correlates with existence of aggressive cancer cells, that dictate the survival of the patient. Stabilisation of prolargin levels in the tumor has the potential of opening a therapeutic window for targeting those cells. High prolargin levels are anti-angiogenic and HCC cancer cells are particularly needy of angiogenesis. With this in mind, preventing prolargin degradation via MMP inhibition is one of the options that could be explored. However, MMP targeting as a treatment concept has suffered from clinical failure because of toxicity, and new strategies need to be developed to eventually bring back MMP inhibitors (MMPi) to the clinic.^42^ In the present study, we had at our disposal only commercially available MMPi that have new generation compounds with better solubility and activity. Indeed, new clinical trials are conducted with more selective MMPi, based also on monoclonal antibodies that specifically target individual MMPs.^43^ Even so, our *in viv*o data showed a remarkable control of tumor growth when sorafenib and batimastat were combined together. While immune-checkpoint inhibitor therapies are bound to change the landscape of HCC treatment, tumor angiogenesis will remain a dominant tumor driver in HCC. The latter will require better treatments and prolargin can – as shown by the present data – assume this role. Further studies will however be necessary to adequately explore how prolargin can be used in synergy with other treatments directed against complimentary microenvironment components. These should involve syngeneic mouse models of HCC where the immune component should be fully exploited.

## Acknowledgement

The authors acknowledge the experimental support of Mrs. Naima Maloujahmoum (Metastasis Research Laboratory, University of Liege), Mrs. Evgenia Turtoi (Tumor Microenvironment and Resistance to Treatment Lab, IRCM, Montpellier), Dr. Susumu Rokudai (Department of Molecular Pharmacology and Oncology, Gunma University) and Mr. Tadashi Handa (Pathology Dept. Gunma University). The authors are particularly thankful to Dr. Arnaud Blomme (Metastasis Research Laboratory, University of Liege) and Mrs. Touko Hirano (Laboratory for Analytical Instruments, Gunma University Graduate School of Medicine) for the help concerning the MS analysis of patient material. The authors thank the Small Animal Imaging Platform of Montpellier (IPAM, http://www.ipam.cnrs.fr/) for the help with animal experiments. AT is thankful to Prof. Jacques Colinge (Bioinformatics and Systems Biology group, IRCM, Montpellier) for his R teachings and to Prof. Peter Friedl (The University of Texas MD Anderson Cancer Center, Houston, Texas, USA) for the helpful discussions.

## Abbreviations used in this paper

HCC: hepatocellular carcinoma
CAF: cancer-associated fibroblasts
EMT: epithelial-mesenchymal transition
PRELP: prolargin
FFPE: formalin-fixed paraffin-embedded
NLF: normal liver fibroblast
CM: conditioned medium
TFA: trifluoroacetic acid
SLRP: small leucine-rich proteoglycan
MMP1: matrix metalloproteinase-1
MMP3: matrix metalloproteinase-3
PDAC: pancreatic ductal adenocarcinoma
Alex: Alexander hepatoma cell line.

## Notes

**Funding sources**: This work was supported with grants from the University of Liège, National Fund for Scientific Research (FNRS), Gunma University (GIAR Research Program for Omics- Based Medical Science). BC is supported by a Fondation de France grant (No. 00078461). RR is supported by Associazione Italiana per la Ricerca sul Cancro (AIRC), grant number MFAG 18459 and IG 2019 - ID. 23151; SR is supported by Fondazione Umberto Veronesi fellowship. AT is a senior research fellow of the French National Institute of Health and Medical Research (INSERM) and is supported by LabEx MabImprove Starting Grant. OD is supported by a grant from the “Fondation Contre le Cancer”. No funding bodies had any role in study design, data collection and analysis, decision to publish, or preparation of the manuscript.

### Competing Interest Statement

The authors have declared no competing interest.

### Summary of Updates

Updated version to consider new developments in systemic therapy of HCC patients. Updated figures 1 and S2.

