## Supplementary Materials, Methods and Figures for "Tumor-Antagonizing Fibroblasts Secrete Prolargin as Tumor Suppressor in Hepatocellular Carcinoma"

### Supplemental Experimental Procedures

#### *Patient information and clinical samples*

HCC samples were obtained from the institutional biobank of the University Hospital Liège, Belgium and Gunma University Hospital, Japan. According to Belgian law, patients were informed that the residual material from surgical procedures could be used for research purposes, and consent is presumed as long as the patient does not oppose (opt out). All Japanese patients provided written informed consent before registration. A total of 188 HCC patients were enrolled in the present study. Patients' survival data was obtained from the individual medical records and preserved strictly by administrators who were independent from the study. Tumor specimens were stored in liquid nitrogen tank before use.

#### *Proteomic Analysis of Accessible Proteins*

Six freshly collected human hepatocarcinoma biopsies and normal liver counterparts were obtained from the University Hospital of Liège, Belgium. Samples were processed according to ex vivo biotinylation procedure as described elsewhere.[1] Briefly, biopsies were sliced in small pieces and soaked in freshly prepared EZ-link Sulfo NHS-SS biotin (1 mg/mL in PBS (phosphate buffered saline), Pierce, Rockford, IL, USA) solution. After 20 min incubation at 37 °C, the reaction was stopped by adding Tris-HCL, pH 7.4 at final concentration of 50 mM. Samples were snap frozen, pulverized and further subjected to affinity purification of biotinylated proteins using streptavidin. Five µg of purified proteins were digested with trypsin (catalog no. 29341524, Promega, Madison, WI, USA) using a 1:50 enzyme to protein ratio. Peptides were further desalted using ZipTip (catalog no. ZTC18S096, Merck, Darmstadt, Germany) according with the manufacturers protocol and analysed using 2D-LC-MS/MS analysis. Three microgram of peptide solution were injected on the 2D-nanoAquity UPLC system (Waters, Corp., Milford, USA) that was coupled online with a Q-Exactive mass spectrometer (Thermo Scientific, Waltham, MA, USA), operated in nano-electrospray positive ion mode. The analytical conditions used were described elsewhere.[2] Raw MS files were analyzed with MaxQuant Software (version 1.5.2.8, Max Planck Institute of Biochemistry, Martinsried, Germany) as previously outlined.[2] Information on protein subcellular localization and biological functions were determined using gene ontology annotation (GO annotation) available on Uniprot (www.uniprot.org), STRING version 10 (www.string-db.org). Gene expression level of identified proteins in healthy human tissues was assessed consulting BioGPS public mRNA database (www.biogps.org). Multi Experiment Viewer software version 4.8 [3] was used to construct heatmaps both from LFQ values of identified proteins and publicly available gene expression data.

***Immunofluorescence***

Deparaffinization, antigen retrieval and blocking steps are described in the main manuscript (Materials and Methods). The sections were then incubated with the 1st primary antibody anti-PRELP (1:1000) for 2 h at room temperature. The sections were rinsed in TNT buffer (2.4 g Tris-base, 8.8 g NaCl, dissolved in 1L $H_2O$, pH 7.6, with 0.01% Tween-20) three times for 5 min each and then incubated for 30 min at room temperature with corresponding HRP-conjugated secondary antibody (Histofine® Simple Stain™ MAX PO (MULTI), catalog no. 414152F; Nichirei Biosciences Inc., Tokyo, Japan). After washing with TNT buffer (three times for 5 min each), fluorescein reagent was applied to the tissue section for 10 min. Following another step of antigen retrieval and blocking, the same process was repeated for anti-alpha-SMA (1:1500, catalog no. M0851; Dako) and anti-CD45 (1:3, clone 2B11 + PD7/26, catalog no. GA751; Dako). All sections were counterstained using DAPI and examined under an All-in-One BZ-X710 fluorescence microscope.

***Immunohistochemistry***

Deparaffinization, antigen retrieval and blocking steps have been performed as described in the main manuscript (Materials and Methods). The following antibodies were used: anti-PRELP (dilution 1:100, catalog no. AF6447 R&D Systems, Minneapolis, MN, USA), anti-MMP3 (dilution 1:1000, catalog no. 500-P324; PeproTech) and anti-CD31 (clone JC70A, catalog no. M0823, Dako). The incubation with the primary antibody was performed overnight at 4°C. Following this, the slides were washed with PBS for 10 min. Signal detection was performed with DAB as described in the main manuscript (Materials and Methods).

Scoring of protein expression was performed in accordance with the previously published methodology.[4] Briefly, each IHC slide was assessed for the intensity of the staining using the following scale: 0 = no staining, 1 = weak, 2 = moderate and 3 = strong. The tissue was further evaluated for the extent of positivity (percent positive area) using the following scale: 0 = 0%–25%, 1 = 25%–50%, 2 = 50%–75% and 3 = 75%–100%. The values obtained by each of the two scales were multiplied to yield a composite value called the IHC score. Pictures of representative fields were taken under a Leica DMRB light microscope (Leica, Wetzlar, Germany). Two independent pathologists scored the samples using the above method and average scores were reported.

***Cell culture***

Human hepatocellular carcinoma (HCC) cell lines HepG2, HUH7, Alexander (PLC/PRF/5), HLE and HLF were obtained from Japan Collection of Research Bioresources Cell Bank (Osaka, Japan). H-6019 human primary liver fibroblast cell line (NLF) was obtained from Cell Biologics

(Chicago, IL, US). Both cancer cells and liver fibroblasts were maintained in DMEM supplemented with 10% heat inactivated FBS and 1% penicillin/streptomycin (all from Gibco, Thermo Fisher Sci., Waltham, MA, USA).

HLF cancer cells were silenced for MMP1 and MMP3 expression using following siRNA (Horizon Discovery, Waterbeach, UK): human anti-*MMP1* (siGENOME Human MMP1, catalog no. M-005951-01-0005), anti-*MMP3* (siGENOME Human MMP3, catalog no. M-005968-03-0005), scramble siRNA (ON-TARGETplus NonTargeting Control Pool, catalog no. D-001810-10-05). The cells were transfected with 20 nM of each siRNA using Lipofectamine (Lipofectamine 2000 reagent, catalog no. 11668-019, Life Technologies, Carlsbad, CA, USA).

Human umbilical vein endothelial cells (HUVEC) were isolated from umbilical cords. They were used at early passages (passages II–V) and grown on plastic surface coated with porcine gelatin in M199 medium (Invitrogen) supplemented with 20% FCS, 100 µg/mL endothelial cell growth factors (Sigma-Aldrich) and 100 µg/mL porcine heparin (Sigma-Aldrich). Sub-confluent cultures (70%–90% confluence) of low passages (until passage 8) were utilized for all experiments. Conditioned medium (CM) from hepatocellular carcinoma cell lines was obtained following 48h of incubation of 80% confluent cells in serum-free medium. Cells CM were collected, centrifuged for 5 min at 150xg, room temperature, and then added to fibroblast monolayer (cells were pre-starved in serum-free medium for 6h) for an additional 48h. Following this, CM were collected, centrifuged for 5 min at 150xg at room temperature, and then further used for WB, ELISA or proteomic analysis. For denaturing experiment, CM collected was boiled 10 minutes at 99 °C, or frozen and thawed twice. Fibroblast monolayers were washed two times with PBS and then either lysed for Western blot analysis or used for RNA extraction.

***In vitro colony formation assay***

For colony formation, HCC cells were seeded into 6-well plate at a density of 1000 cells/well. After 8 days of culture, cells were fixed and stained with a solution of 0.1% crystal violet and the colonies containing over 20 cells were counted under the microscope visually. Experiment were done in triplicates and repeated three times.

***Western blot analysis***

Total proteins from both tissues and cells were extracted using 1% SDS buffer [40mM Tris-HCl (pH 7.6), 1% sodium dodecyl sulfate (SDS) and protease/phosphatase inhibitor cocktails (catalog no. 16829900; Sigma-Aldrich). CM samples were concentrated 10-fold using Amicon Ultra-0.5 3kDa filters (catalog no. UFC500324; Millipore), while the cell culture medium was exchanged with the lysis buffer (same as described above). The protein content of all samples was determined using the Pierce BCA Protein Assay Kit (catalog no. 23225; Thermo Scientific).

Twenty micrograms of proteins were supplemented with Laemmli buffer (0.1% 2-mercaptoethanol, 0.0005% bromophenol blue, 10% glycerol, 2% SDS in 63 mM Tris-HCl (pH 6.8)), were separated on 10% polyacrylamide denaturing gel and transferred to nitrocellulose membranes for Western blotting. Following antibodies were used for WB analysis: anti-PRELP (dilution 1:500, catalog no. AF6447; R&D Systems), anti-pMET (Tyr1234/1235) (dilution 1:1000, catalog no. 3077; Cell Signaling), anti-MET (dilution 1:1000, catalog no. 8198, Cell Signaling), anti-TUBB (dilution 1:1000, catalog no. 2128, Cell Signaling), anti-HSC70 mAb (dilution 1:1000, catalog no. sc-7298; Santa Cruz Biotechnology) and anti-ACTB (dilution 1:1000, catalog no. 3700, Cell Signaling). Cell extracts were normalized using anti-HSC70, -ACTB or –TUBB. Conditioned media were normalized for protein load by staining with SYPRO™ Ruby Protein Blot Stain (Thermo Scientific; catalog no. S11791).

***Evaluation of PRELP degradation***

Five hundred microliters of CM from HUH7 or HLF cells were pre-incubated with 10 µM amiloride hydrochloride hydrate (catalog no. A7410; Sigma), 100 nM batimastat, (catalog no. SML0041; Sigma), 100 nM GM6001 (Sigma, catalog no. CC1000, Saint Louis, MI) or protease inhibitors 1X (Roche Molecular Systems, protease inhibitor cocktail tablets, catalog no. 16829900) for 1h at 37 ºC under agitation. Following this, 100 ng recombinant PRELP (catalog no. 6447-PR; R&D Systems) was added and incubated at 37 ºC under agitation for 30 minutes, and afterwards for 2, 4, 6, and 24 hours respectively. PRELP degradation was evaluated using WB analysis.

***Gene expression analysis***

Total RNA was isolated with the Nucleospin RNA Isolation Kit (catalog no. MN740955; Macherey-Nagel, Dueren, Germany). According to the manufacturer's instructions, one microgram of RNA was reverse-transcribed using the SuperScript III Reverse Transcriptase (catalog no. 18080; Invitrogen, Carlsbad, CA, USA). Twenty nanograms of cDNA were used for respective PCR reactions. cDNA was mixed with primers (0.5 µM), 0.2 ul human UPL-probe system (Roche, Mannheim, Germany) (#34 for *PRELP*, catalog no. 0487671001; #7 for matrix metalloproteinase-1 (*MMP1*), cat. no. 04685059001; #58 for metalloproteinase-3 (*MMP3*), cat.no. 04688554001, #64 for beta-actin (*ACTB*), cat. no. 04688635001) and Kapa Probe Fast qPCR kit Master Mix (2X) (catalog no. KK4702; Sigma Aldrich, St. Louis, MO, USA). Quantitative real-time PCR (qRT-PCR) was performed using the LightCycler 480 system (Roche) and the corresponding manufacturer software. The following cycling conditions were used: 95°C for 3 min then 40 cycles of 95°C (3 sec) and 60°C (30 sec). Sequences of each primers were as follows: *PRELP*, forward 5'-GCT CAA AGA GGC CGA GAA A-3' and reverse 5'-AGC AGC TTT GCC AGA AGG -3'; *MMP1*, forward 5'-GCT AAC CTT TGA TGC TAT AAC TAC GA-3'; and reverse 5'-TTT GTG CGC ATG TAG AAT CTG; *MMP3,* forward 5'-GCA GTT TGC TCA GCC TAT CC-3'; and reverse 5'-TTT CTC CTA ACA AAC TGT TTC ACA TC-3'; *18S,* forward 5'-CTT CCA CAG GAG GCC TAC AC-3'; and reverse 5'-CGC AAA ATA TGC TGG AAC TTT-3'. The relative gene expression levels were normalized using 18S RNA.

***PRELP silenced stable clones***

Stable PRELP depletion was achieved in NLF using lentiviral shRNA particles. Prolargin shRNA (TRCN0000162750) cloned into pLKO.1 Puro, Sigma, St. Louis, MO, USA) or control shRNA (Cat. #: SHC005; Sigma, St. Louis, MO, USA) were cotransfected with GAG/POL (Addgene, Cambridge, MA, USA) and a VSV-G encoding plasmids (Addgene, Cambridge, MA, USA) in lenti-X 293T cells (Clontech, Mountain view, CA, USA). Forty-eight hours post-transfection of 293T cells, viral supernatants were collected, filtrated, and concentrated at 1,500×g by ultracentrifugation. After transduction, the stably clones were selected with puromycin treatment (1 µg/mL).

***Chick chorioallantoic membrane model***

Three days post-fertilization, eggs were fenestrated and 8ml of albumin was removed in order to expose the chorioallantoic membrane (CAM). Eggs were sealed with Durapore tape (3M, Diagem, Belgium) and kept closed at 37°C (80% humidity) until implantation of the cells. Four million HLF cells and $1x10^6$ NLF cells suspended in 20µL of culture medium (without serum) were mixed with 20 µL of matrigel and were co-implanted on the CAM. In order to test the efficacy of batimastat and sorafenib combination treatment, 4 different conditions have been established: CTR, for placebo (n=5), batimastat alone (n=5), sorafenib alone (n=5) and batimastat + sorafenib (n=5). CAM were treated three times during the tumor developing. Batimastat and Sorafenib have been used both at 50 nM final concentration. Tumors were allowed to develop for an additional 7 days and at day 18 post-fertilization, the tumors were collected and the volume was calculated using the formula as indicated above.

***Murine in vivo models***

Additionally to ARRIVE guidelines, animal experimentation adhered to the Guide for the Care and Use of Laboratory Animals prepared by the Institute of Laboratory Animal Resources of the National Research Council and published by National Academies Press, as well as to European and local legislation. In France, mice were obtained from Charles River and were kept in pathogen-free conditions in the animal facilities of the Institut de Recherche sur le Cancer de Montpellier. In Japan, mice were purchased from CLEA Japan and housed in the Bioresource Center in Gunma

University (12h light/dark cycle, lights on at 6 a.m.). They were acclimated to the room 1 week before the beginning of the following experiment. Food and water were provided ad libitum. HLF-RFP-Luc cells ($1.5x10^6$ cells) were suspended in 20 µl of cell culture medium and growth-factor-reduced Matrigel 1:1 and injected into the liver of six-week old athymic nude mice. One-week post implantation mice were randomly divided in two groups (each n=5). One group of mice was treated with sorafenib 30 mg/kg alone (catalog no. 1009644; Cayman Chemical Company, Michigan, US) or in combination with batimastat (catalog no. SML0041, Sigma) 15 mg/kg during 5 weeks. Tumor growth in both experiments was monitored using bioluminescence on an IVIS Lumina II Imaging System (Perkin Elmer, Waltham, MA, USA) after luciferine (catalog no. 122796; Perkin Elmer) IP injection of 200 µL of 10 mg/ml luciferine solution in PBS with signal acquisition at 10 min post injection. All animals were sacrificed at day 52.

***Proteomic analysis of cellular secretome & PRELP interactome***

Concentrated conditioned media (corresponding to 5 µg of proteins) from different cancer cell lines were lyophilized and re-suspended in 50 µL of 100mM ammonium bicarbonate solution, containing 0.1% RapiGest SF (catalog no.: 186001861; Waters Corporation, Milford, MA, USA). Proteins were further reduced in 1,4-dithiothreitol (10 mM) (catalog no.: D0632-10G; Sigma-Aldrich) for 30 minutes at 60°C and then alkylated using 2-chloroacetamide (22mM) (catalog no.: 30208220; Sigma-Aldrich) for 30 minutes at RT and in darkness. Protein digestion was performed overnight at 37°C with trypsin (catalog no.: 29341524; Promega, Madison, WI, USA) using 1:50 enzyme to protein ratio. Following the digestion, 1µL of PNGase enzyme (catalog no.: P0704S; BioLabs, MA, USA) was added to each preparation and samples were incubated at 37°C for further 4h. Peptides were acidified with 0.5% (final) TFA, incubated for 1h at 37°C and then centrifuged at 13000xg for 10 minutes. The supernatant containing peptides was desalted using ZipTip (catalog no.: ZTC18S096; Merck, Darmstadt, Germany) according with the manufacturer's protocol.

For PRELP interactome analysis, 300 ng of recombinant PRELP was incubated for 30min at room temperature with 300 µL of concentrated conditioned media (corresponding to 10 µg of proteins) from HLF and HUH7 cells respectively. Following this 3 µg of PRELP antibody (same clone as used in WB and IHC analysis) was added to the mixture and the samples were further incubated overnight at 4°C (under rotation). Following overnight incubation (16h), 5 µl of Protein G magnetic beads (catalog no.: G747A; Promega, Madison, WI, USA) (pre-washed with PBS) were added to the samples. The samples were further incubated at 4°C (under rotation) for additional 4h. Magnetic beads were pulled down using appropriate magnet and then washed 3x with PBS for 5min. Bound proteins were released using 2x 20 µL of 0.5% TFA in water. The collected eluates were pooled, lyophilized and re-suspended in 50 µL of 0.1% RapiGest SF/ 100 mM ammonium bicarbonate solution. Further processing of samples was conducted according to the same procedure as described for the secreted proteins above.

Peptide samples from both experiments outlined above were analyzed using 1D-LC-MS/MS analysis. Briefly, Ekspert nanoLC400 (Eksigent, Dublin, CA, USA) was coupled with TripleTOF-5600 (AB Sciex, Concord, ON, Canada). Five µg of sample was injected on the C18 analytical column (Acclaim PepMap® 75 µm x 150 mm, p/n: 160321; Thermo Fisher) with a gradient of 0–40% phase B (90 % acetonitrile, 10.9 % water and 0.1 % formic acid) for 80min at the flow rate of 0.3 µl/min. Peptide masses were surveyed over a mass range from 400 to 1600 m/z and up-to 30 MS/MS spectra per one survey scan were collected in data dependent fashion. Raw MS files were analyzed with MaxQuant Software as previously outlined.[2] Peptides counts were used for relative quantification.

***Endothelial cells & in vitro activity assays***

Tumor cells were seeded in complete medium at 50,000 cells/cm$^2$. The day after, cells were washed and grown in absence of serum for 48 hours. The conditioned media were collected, filtered and used for the *in vitro* assays. HUVEC proliferation assay: HUVEC were seeded at 15,000 cells/cm$^2$ and then treated with 100% conditioned media in 2.5% FCS in presence or absence of 4 µg/mL PRELP recombinant. Twenty-four hours later, cells were detached and counted using MACS Quant cytofluorimeter (Milteny Biotec). HUVEC sprouting assay: HUVEC spheroid aggregates were first embedded in fibrin gel and then stimulated with 50% of conditioned media, 5% FCS containing growth factors combined (or not) with 4 µg/mL PRELP. After 24 hours, growing cell sprouts were photographed and counted under an inverted microscope. HUVEC wound repair assay: HUVEC monolayers were scratched with a 200 µL tip to obtain a 2-mm-thick denuded area and cultured in the presence of growth factors with or without the addition of 4 µg/mL recombinant PRELP in 3.5% FCS. After 18 hours wounded monolayers were photographed and the percentage of repaired area was quantified with Fiji software.[5]

***Surface plasmon resonance analysis***

Surface plasmon resonance (SPR) measurements were performed on a BIAcore X100 instrument (GE-Healthcare, Chicago, IL, USA). Recombinant PRELP was immobilized on a CM5 sensor chip (GE-Healthcare) with amine-coupling method, allowing the immobilization of 2800 RU resonance units (RU) (56.8 pmol/mm$^2$ of PRELP). Blank immobilization was performed as control. To analyze their direct binding to sensorchip-immobilized PRELP, the different proteins were suspended at 100 nM in 10 mM HEPES, 150 mM NaCl, 3 mM EDTA 0.05% surfactant P20, pH 7.4 (HBS-EP buffer), injected for 3 minutes (to allow their association to PRELP) and washed until dissociation was observed. Following growth factors were tested: fibroblast growth factor 1

(FGF1) and FGF2 (Tecnogen, Caserta, Italy), transforming growth factor-β1 (TGF-β1), placenta derived growth factor 2 (PDFG-2) (ReliaTech, Wolfenbüttel, Germany), hepatocyte growth factor (HGF), connective tissue growth factor (CTGF) and epidermal growth factor (EGF) (PeproTech, Rock Hill, NJ, USA), PRELP, FGF3, bone morphogenetic protein 2 (BMP2), BMP4, interleukin-8 (IL-8), neuroblastoma suppressor of tumorigenicity 1 (NBL1) and tumor necrosis factor α (TNF-α) (R&D Systems), vascular endothelial growth factor-A (VEGF-$A_{165}$ isoform) and VEGF-C were kindly provided by K. Ballmer-Hofer (PSI, Villigen, Switzerland).

***Homology model of PRELP and docking***

PRELP has previously been predicted to possess multiple Leu-rich-Repeats (LRR).[6] Therefore, we aligned the PREPL primary sequence (P51888, residues 75-308) versus the primary sequences of the high-resolution X-ray crystal structures of human fibromodulin (pdb code: 5mx0) and human osteomodulin (pdb code: 5yq5) using PROMALS3D (**Figure S6**).[7] The human PRELP primary sequence shares 55.5% and 54.6% sequence similarity with each structure, respectively. Using the secondary structure-weighted alignment of fibromodulin, osteomodulin, and PRELP, a homology model was computed using Modeller.[8] Both template structures were used in the homology model calculation. Further, the predicted α-helices of the LRR were restrained as α-helices in all of the models. Fifty models were calculated. We selected the lowest scoring model based on the Modeller Objective Function for our subsequent protein docking studies. The initial docking poses of the PRELP homology model and the various growth factors were determined using the ClusPro server.[9] The most likely 25 poses were then scored using FireDock.[10] The pose with the highest Z-score from FireDock was then refined using Rosetta Dock2 in local-refinement mode. Shape complementarity 'Sc' was computed using the 'sc' program available in the CCP4 package.[11]

***Tumor cells & in vitro activity assays***

HLE scattering assay: HLE spheroid aggregates were embedded in fibrin gel and stimulated with 100 ng/mL HGF in the absence or in the presence of 4 µg/mL recombinant PRELP in 5% FCS DMEM. After 24 hours, cell scattering was photographed and counted under an inverted microscope. HLE migration assay: HLE monolayers were scratched with a 200 µL tip to obtain a 2-mm-thick denuded area and cultured in the presence of 100 ng/mL HGF with or without 4 µg/mL recombinant PRELP in 0.4% FCS DMEM. After 18 hours wounded monolayers were photographed and the percentage of repaired area was quantified with Fiji software. To test the activity of HGF/c-MET system, HLE, Alexander and HUVEC cells were seeded 50,000/$cm^2$, serum starved for 5 hours and treated with HGF (100 ng/mL) and PRELP (4 µg/mL). After 15 minutes of incubation cell samples were washed in cold PBS and homogenized in lysis buffer. Samples were quantified and used in Western blot analysis.

***Statistical analysis***

Unless otherwise indicated, statistical analysis was performed using three biological replicates and a two-sided unpaired Student's *t*-test. The *t*-test was used when data followed a normal distribution (Shapiro-Wilk test, threshold 0.05). For IHC evaluation, the number of samples is indicated in the legends to figures. Statistical differences of IHC data were tested with the Mann-Whitney-U-test because these results did not follow the normal distribution (Shapiro-Wilk test, threshold 0.05). Fisher's exact test was used to compare categorical variables for survival analysis. OS was estimated according to the Kaplan–Meier method, and was compared among groups with the log-rank test. All statistical analyses were performed with EZR (Saitama Medical Center, Jichi Medical University, Japan), which is a graphical user interface for R (The R Foundation for Statistical Computing, version 3.3.2). EZR is a modified version of R commander (version 2.3-1) that includes statistical functions that are frequently used in biostatistics.

**Supplemental Results**

**Supplementary Tables**

| | % |
|---|---|
| Gender/Age | |
| Male | 78 |
| Female | 22 |
| Age 60+ | 80 |
| Age 60- | 20 |
| Hepatitis/Cirrhosis | |
| No | 40 |
| B | 6 |
| C | 32 |
| B+C | 6 |
| Chirrosis Viro- | 10 |
| NASH | 4 |
| Not known | 2 |
| pTNM | |
| T1 | 18 |
| T2 | 35 |
| T3 | 26 |
| T4 | 15 |
| Not known | 6 |
| Differentiation | |
| Good | 36 |
| Medium | 25 |
| Poor | 37 |
| Not known | 2 |
| Liver Transplant | |
| Yes | 30 |
| No | 70 |
| Adjuvant Therapy | |
| Yes | 40 |
| No | 60 |
| Surgery | |
| Yes | 100 |
| No | 0 |

**Table S1:** Clinical features of HCC patients used for IHC analysis/survival analysis. 188 HCC patients were involved in the current study.

### Supplementary Figures

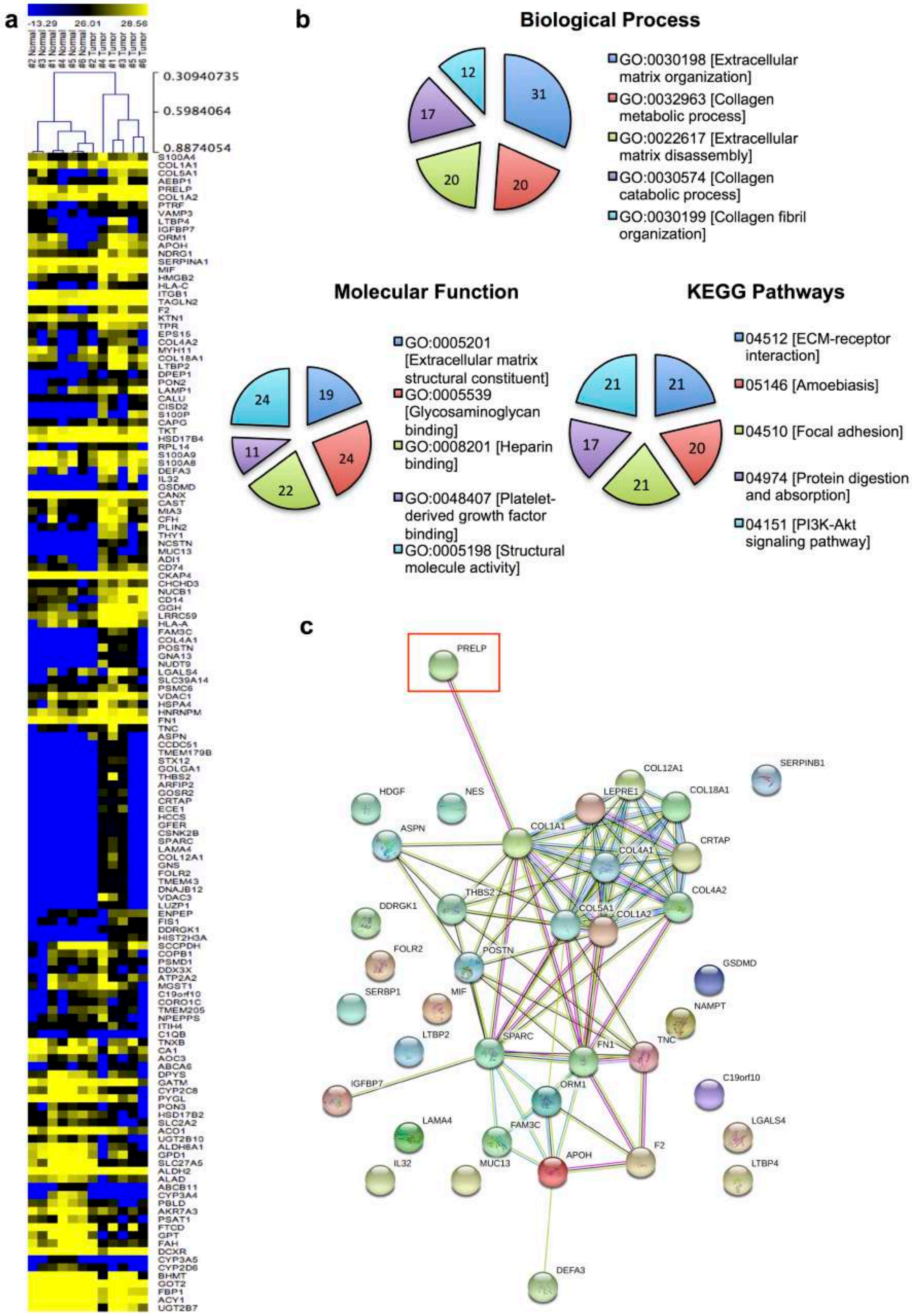

**Figure S1:** MS-based accessibilome analysis of human HCC. **a.** Pearson correlation clustering of proteins identified as modulated (non-tumoral versus tumoral samples) in 3 out of 6 patients. **b.** Classification of modulated proteins according to the GO terms (Gene Ontology) with respect to biological process, molecular function and involvement in pathways (KEGG). The numbers indicated in the pie charts refer to percentage of proteins found in the respective category. **c.** Network analysis using STRING software (as outlined in the Materials and Methods section). Proteins are connected based on the existing data relating to the co-expression, binding & molecular classification of individual proteins.

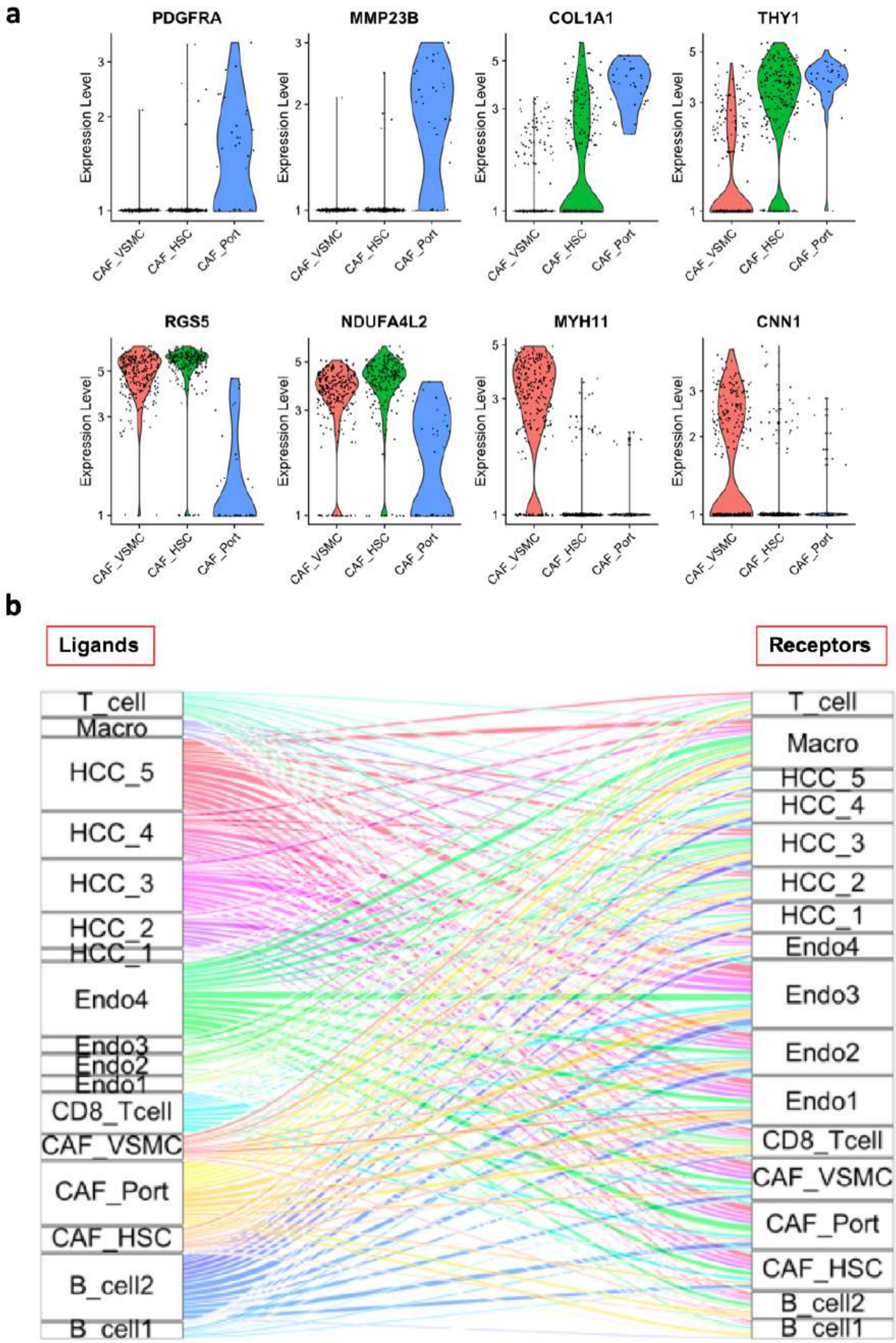

**Figure S2:** Portal-fibroblast derived CAF have strong ligand-receptor communication with other stromal and cancer cells in HCC. **a.** Violin plot of eight major markers delineating the CAF cell of origin in HCC (vascular smooth muscle cell [VSMC], hepatic stellate cell [HSC] or portal fibroblast [Port]). **b.** Ligand-receptor interaction map between stromal and cancer cells in the HCC. Number of interactions is proportional to the size of individual boxes. Panel **a**/**b**: Data set from GEO accession GSE125449.[12]

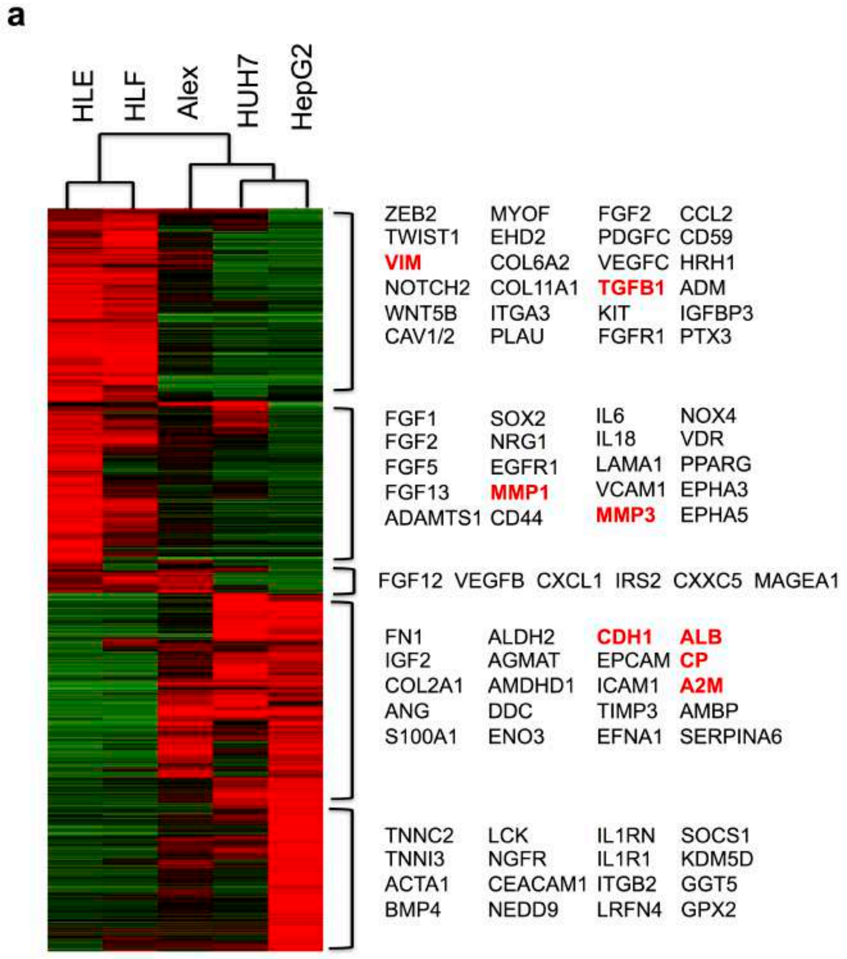

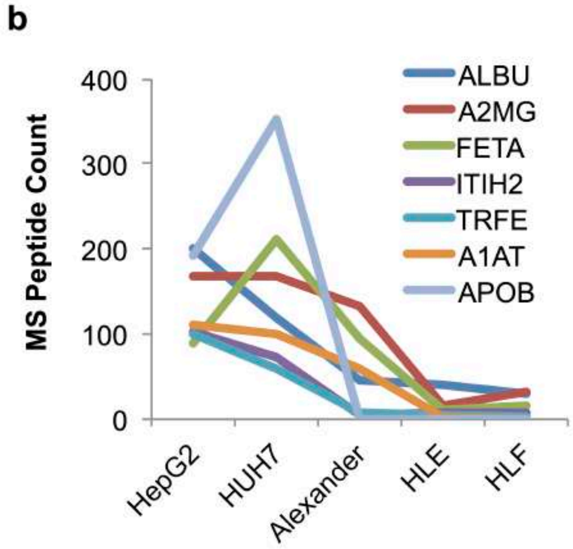

**Figure S3:** Characterization of the HCC cell line panel. **a.** Pearson correlation clustering analysis of publicly available dataset GSE35818.[13] Highlighted are the most well-known genes found in representative clusters. **b.** Proteomic analysis of the secretome obtained from the respective HCC cells. Displayed are the proteins that decrease in expression along with increasing aggressiveness of HCC cells lines.

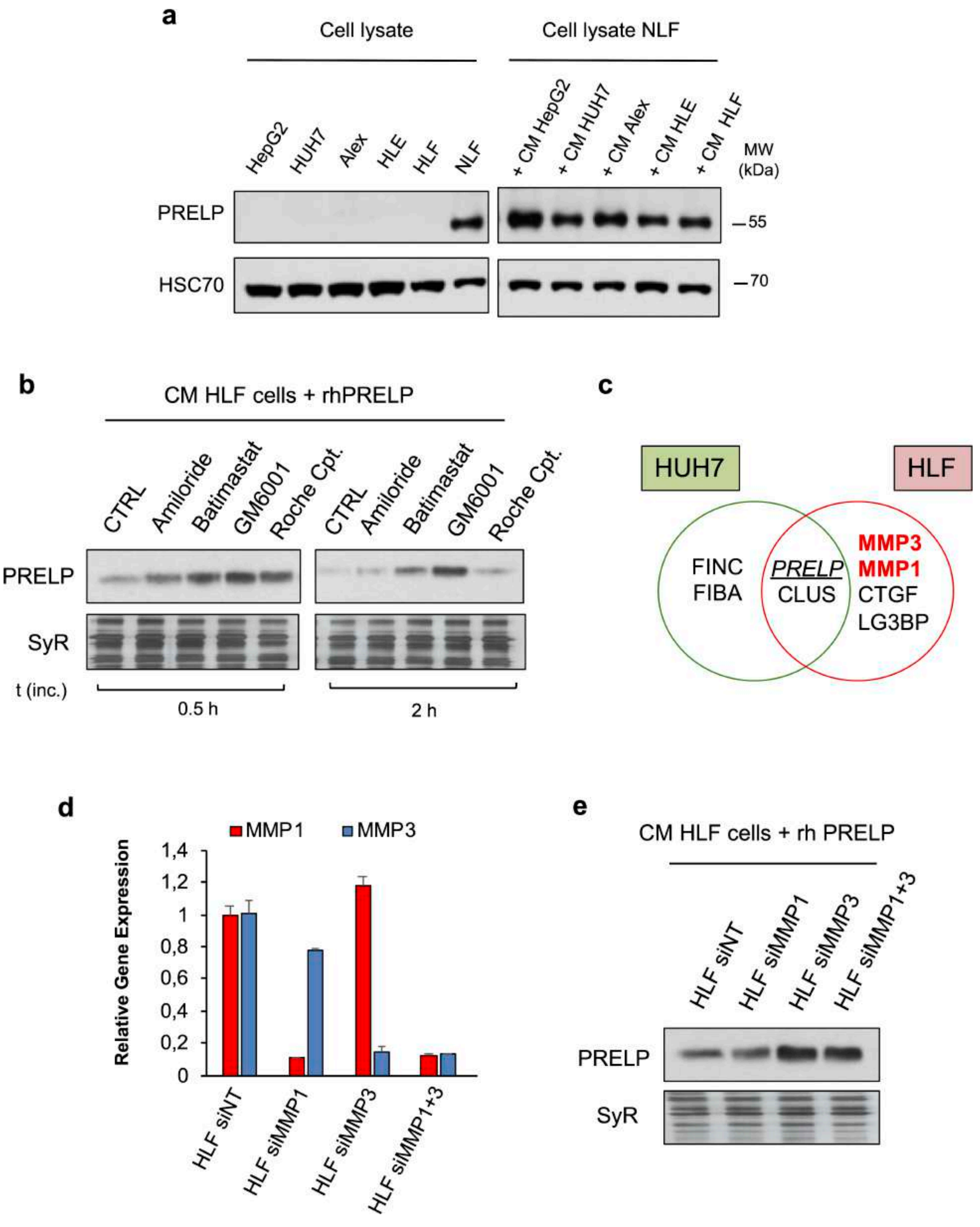

**Figure S4:** MMP1 and MMP3 degrade PRELP. **a.** (*left*) PRELP expression profiles in cell lysates obtained from cancer cells and normal liver fibroblasts (NLF). (*right*) PRELP intracellular levels following the treatment of NLF with cancer cell conditioned media. HSC70 levels were used for normalization. **b.** Treatment of HLF cell-derived conditioned media with various protease inhibitors and their incubation with recombinant prolargin at 37°C. **c.** Prolargin interactome analysis in conditioned media from HUH7 and HLF cells. **d.** RT-qPCR analysis of the efficacy of MMP1 and MMP3 gene expression silencing. **e.** Analysis of prolargin degradation using conditioned media from HLF cells silenced either for MMP1, MMP3 or both. Panels **b** and **e**: total protein stain with SYPRO Ruby (SyR) were used for normalization.

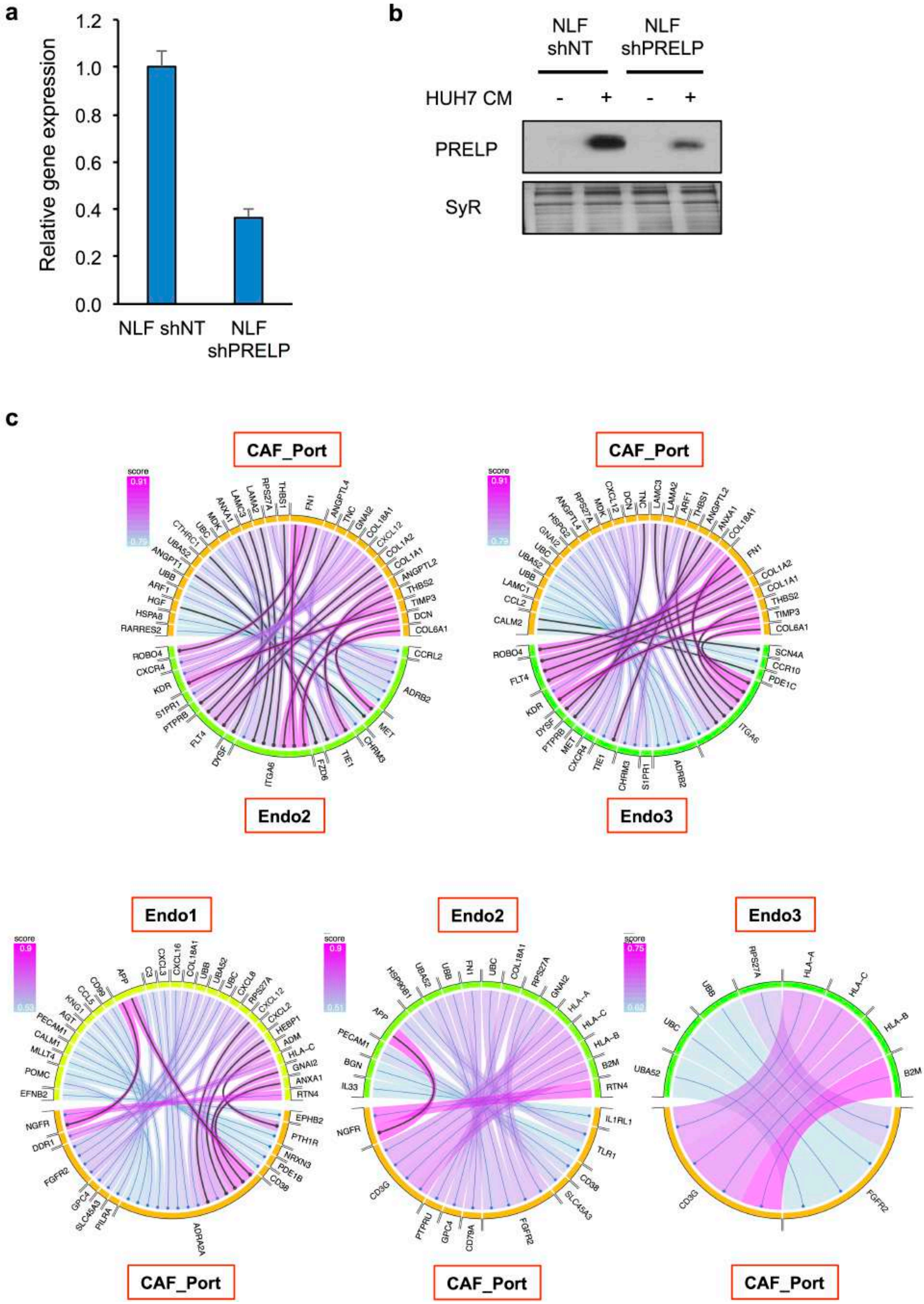

**Figure S5:** Validation of prolargin depleted NLF and ligand-receptor analysis. **a.** PCR analysis of mRNA expression in NLF that have been silenced for PRELP expression. **b.** Western blot analysis of PRELP protein expression in conditioned media of NLF silenced for PRELP and challenged with the CM of HUH7 cancer cells. Panels **a** and **b**: NT denotes mock-transfected cells. **c.** Ligand-receptor analysis of portal-fibroblast derived CAF with different populations of endothelial cells (see **Figure 1a**). Panel **c**: Data set from GEO accession GSE125449.[12]

**a**

```
osteomodulin   SECFCPTNFPSSMYCDNRKLKTIPNIPMHIQQLYLQFNEIEAVTANSFINATHLKEINLS 123
fibromodulin   QECDCPPNFPTAMYCDNRNLKYLPFVPSRMKYVYFQNNQITSIQEGVFDNATGLLWIALH 137
prolargin      RECYCPPDFPSALYCDSRNLRKVPVIPPRIHYLYLQNNFITELPVESFQNATGLRWINLD 134

osteomodulin   HNKIKSQKIDYGVFAKLPNLLQLHLEHNNLEEFPFPLPKSLERLLLGYNEISKLQTNAMD 183
fibromodulin   GNQITSDKVGRKVFSKLRHLERLYLDHNNLTRMPGPLPRSLRELHLDHNQISRVPNNALE 197
prolargin      NNRIRKI--DQRVLEKLPGLVFLYMEKNQLEEVPSALPRNLEQLRLSQNHISRIPPGVFS 192

osteomodulin   GLVNLTMLDLCYNYLHDSLLKDKIFAKMEKLMQLNLCSNRLESMPPGLPSSLMYLSLENN 255
fibromodulin   GLENLTALYLQHNEIQEV---GSSMRGLRSLILLDLSYNHLRKVPDGLPSALEQLYMEHN 254
prolargin      KLENLLLLDLQHNRLSDGVFKPDTFHGLKNLMQLNLAHNILRKMPPRVPTAIHQLYLDSN 252

osteomodulin   SISSIPEKYFDKLPKLHTLRMSHNKLQD--IPYNIFNLPNIVELSVGHNKLKQAFYIPRN 301
fibromodulin   NVYTVPDSYFRGAPKLLYVRLSHNSLTNNGLASNTFNSSSLLELDLSYNQLQKIPPVNTN 315
prolargin      KIETIPNGYFKSFPNLAFIRLNYNKLTDRGLPKNSFNISNLLVLHLSHNRISSVPAINNR 312

osteomodulin   LEHLYLQNNEIEKMNLTVMCPS---------IDPLHYHHLTYIRVDQNKLKE-PISSYIF 351
fibromodulin   LENLYLQGNRINEFSISSFCTV---------VDVVNFSKLQVLRLDGNEIKRSAMPADAP 365
prolargin      LEHLYLNNNSIEKINGTQICPNDLVAFHDFSSDLENVPHLRYLRLDGNYLKP-PIPLDLM 371

osteomodulin   FCFPHIHTIYYGEQ 365
fibromodulin   LCLRLASLIEI--- 376
prolargin      MCFRLLQSVVI--- 382
```

**b**

**c**

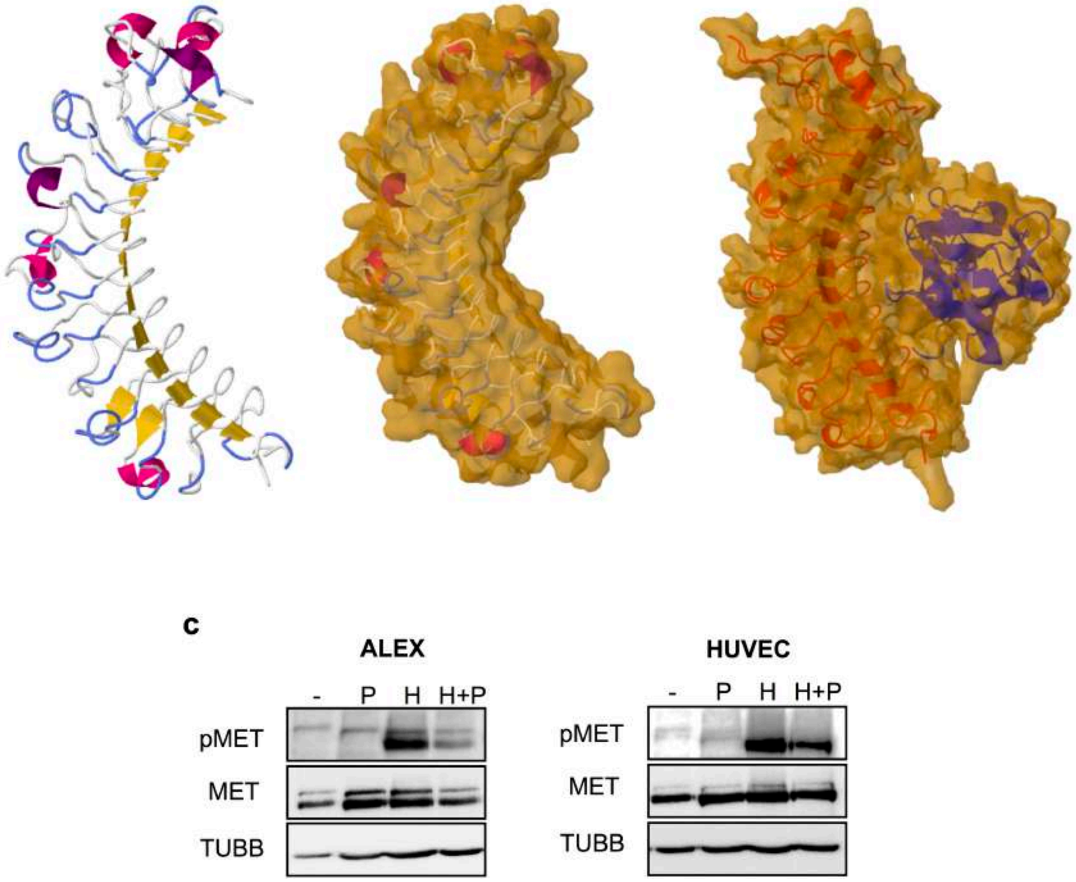

**Figure S6:** PRELP structure and growth factor binding activity. **a.** Primary sequence alignment of the template structures used to compute the homology model of prolargin. The 3D structures of Osteomodulin (Q99983, pdb:5YQ5) and Fibromodulin (Q06828P, pdb: 5MX1) were aligned with the human PRELP sequence (P51888) using PROMALS3D.[7] The red letters in the PRELP sequence correspond to the 11-residue consensus sequence, LxxLxLxxNxL, typical of Leucine-Rich-Repeats (LRR) proteins. **b.** Inferred PRELP 3D structure (backbone and with calculated molecular surface) alone or docked with FGF2. **c.** Analysis of PRELP binding activity to HGF using Western blot. Evaluated is activation of MET receptor in Alexander and HUVEC cells. Following abbreviations were used: prolargin (P), HFG (H) and mix of prolargin and HGF (H+P). TUBB is used as loading control.
